# Identification of SLC35A1 as an essential host factor for the transduction of multi-serotype recombinant adeno-associated virus (AAV) vectors

**DOI:** 10.1101/2024.10.16.618764

**Authors:** Xiujuan Zhang, Siyuan Hao, Zehua Feng, Kang Ning, Cagla Aksu Kuz, Shane McFarlin, Donovan Richart, Fang Cheng, Ander Zhang-Chen, Richenda McFarlane, Ziying Yan, Jianming Qiu

## Abstract

We conducted a genome-wide CRISPR/Cas9 screen in suspension 293-F cells transduced with rAAV5. The highly selected genes revealed after two rounds of screens included the previously reported *KIAA039L*, *TM9SF2*, and *RNF121*, along with a cluster of genes involved in glycan biogenesis, Golgi apparatus localization and endoplasmic reticulum penetration. In this report, we focused on solute carrier family 35 member A1 (*SLC35A1*), a Golgi apparatus-localized cytidine 5’-monophosphate-sialic acid (CMP-SIA) transporter. We confirmed that *SLC35A1* knockout (KO) significantly decreased rAAV5 transduction to a level lower than that observed in *KIAA0319L* or *TM9SF2* KO cells. Although *SLC35A1* KO drastically reduced the expression of α2,6-linked SIA on the cell surface, the expression of α2,3-linked SIA, as well as the cell binding and internalization of rAAV5, were only moderately affected. Moreover, *SLC35A1* KO significantly diminished the transduction of AAV multi-serotypes, including rAAV2 and rAAV3 which do not utilize SIAs for primary attachment. Notably, the SLC35A1 KO markedly increased transduction of rAAV9 and rAAV11, which primarily attach to cells via binding to galactose. Further analyses revealed that *SLC35A1* KO significantly decreased vector nuclear import. More importantly, although the C-terminal cytoplasmic tail deletion (ΔC Tail) mutant of SLC35A1 did not drastically decrease SIA expression, it significantly decreased rAAV transduction, as well as vector nuclear import, suggesting the C-tail is critical in these processes. Furthermore, the T128A mutant significantly decreased SIA expression, but still supported rAAV transduction and nuclear import. These findings highlight the involvement of the CMP-SIA transporter in the intracellular trafficking of rAAV vectors post-internalization.

**IMPORTANCE:** rAAV is an essential tool for gene delivery in the treatment of genetic disorders, yet the mechanisms of rAAV transduction remain partially understood. GPR108 is vital for the transduction of most rAAV vectors, but not for rAAV5. We aimed to identify host factors that impact AAV5 transduction akin to GPR108. Using a genome-wide CRISPR/Cas9 screen in 293-F cells, we identified SLC35A1, a Golgi apparatus-localized CMP-sialic acid transporter that transports CMP-sialic acid from cytoplasm into the Golgi apparatus for sialylation, is essential to rAAV transduction. Further studies across various AAV serotypes showed SLC35A1 significantly affects vector nuclear import post-internalization. These results underscore the crucial role of SLC35A1 in intracellular trafficking beyond the initial cell attachment of rAAV.

## INTRODUCTION

Recombinant adeno-associated viruses (rAAVs) are powerful vectors for gene therapy, offering significant advantages due to their ability to transduce a wide variety of cell types, non-pathogenic nature, and potential for long-term persistence of therapeutic gene expression (1,2). In the past few years, gene therapy has had great success with six rAAV-based medicines approved by the US FDA (3–7), and more than 350 AAV-based gene therapies undergoing clinical trials worldwide. Among the multiple AAV serotypes, AAV5 has shown distinct properties that make it particularly effective for targeting liver, airway epithelia, vascular endothelial cells, and smooth muscles (8). rAAV5-based gene therapies, ROCTAVIAN and HEMGENIX, for hemophilia A and B, respectively, are currently employed clinically (5,7). Importantly, an AAV5 variant, AAV2.5T, was developed by directed evolution of the capsid gene, shows a high tropism to human airways (9), and has demonstrated a functional correction in the treatment of cystic fibrosis (CF) in an *in vitro* CF airway epithelial model and a preclinical trial (10,11). The rAAV2.5T capsid is a chimera of the VP1-unique domain (aa1-119) (VP1u) of AAV2 with the remainder (aa120-725) of the AAV5 capsid, along with a key point mutation (A581T) of AAV5 VP1 (9).

The efficiency of AAV-mediated gene delivery relies on its ability to enter the host cell and navigate through intracellular compartments to reach the nucleus. AAV entry is initiated by the attachment to specific cell surface glycan(s) and further requires a proteinaceous receptor, which determines the tissue tropism of different capsids (12–14). A variety of glycans have been identified as the attachment receptors used by AAV vectors (15). In general, AAV serotype vectors can be grouped into 3 categories concerning their glycan receptor usage: heparan sulfate proteoglycan (HSPG) for AAV2, AAV3, and AAV13 (16–18); α2,3-and α2,6-linked sialic acid (SIA) for AAV1, AAV4, AAV5, and AAV6; terminal N-linked galactose for AAV9 (19,20). Among SIA-used AAVs, both α2,3 and α2,6 N-linked SIAs are used for AAV1 (21) and AAV6 (21,22); α2,3 O-linked SIA for AAV4 (23), α2,3 N-linked SIA for AAV5 (23,24) and AAV2.5T (25). A transmembrane protein KIAA0319L serves as a multi-serotype AAV receptor (AAVR) (26,27), but not for AAV4 and its related serotypes (28).

After internalization via endocytosis, AAV traffics to the *trans*-Golgi network (TGN) through various endosomes and/or the syntaxin 5-positive (STX5^+^) vesicle (29,30). During this process, the acidic milieu within these membrane vesicular compartments induces a conformational change in the AAV capsid, leading to the extrusion of the VP1u from the capsid surface (31–33). VP1u contains a phospholipase A2 (PLA_2_)-like activity domain that is essential for AAV to escape from these vesicles for nuclear entry (34). Once inside the nucleus, AAV undergoes uncoating to expose its single-stranded (ss)DNA genome, which is then converted to transcription-competent double-stranded (ds)DNA (1).

Using a genome-wide gene knockout screen, several host cell factors that restrict AAV entry and intracellular trafficking have been identified, including KIAA0319L (AAVR), G protein-coupled receptor 108 (GPR108), ring finger protein 121 (RNF121), and WD repeat domain 63 (WDR63), for various AAV serotypes (26,35–38). While KIAA0319L serves as a proteinaceous receptor for virion entry (26), GPR108, a protein localized to the *trans*-Golgi network (TGN), mediates the post-entry trafficking of several AAV serotypes through interacting with VP1u (35). However, *GPR108* knockout (KO) does not affect rAAV5 transduction (35), suggesting that other host factors are involved in loading AAV5 to the TGN.

In this study, we employed a genome-wide CRISPR/Cas9 screening approach in 293-F cells to identify host factors that affect rAAV5 transduction. Our objective was to uncover genes whose disruption impedes AAV5 entry, intracellular trafficking, or transgene expression in a selected cell population resistant to rAAV5 transduction. The screen successfully identified several known factors, including *KIAA0319L*, *TM9SF2*, and *RNF121*, as well as novel candidates involved in glycan biogenesis and endoplasmic reticulum (ER) penetration. Among these, solute carrier family 35 A1 (SLC35A1) emerged as a significant player in rAAV5 transduction. SLC35A1, a cytidine 5’-monophosphate-sialic acid (CMP-SIA) transporter localized in the Golgi apparatus, plays a critical role in glycan biogenesis by transporting CMP-SIA from the cytoplasm into the lumen of Golgi apparatus (39). This function is essential for the proper sialylation of glycoproteins and glycolipids, the processes that can influence various cellular activities, including its permissibility for viral infection (40). The structure of SLC35A1 includes specific domains essential for its transport activity (41).

We further validated the role of SLC35A1 in the transduction of other serotypes of AAV, including the AAV5 airway-tropic variant rAAV2.5T, and investigated the mechanisms underlying SLC35A1-mediated AAV transduction. Our study revealed that SLC35A1 is critical for AAV post-entry intracellular trafficking. These findings provide new insights into the role of CMP-SIA transporter in facilitating efficient nuclear import of multi-serotype AAVs. This discovery offers potential avenues for optimizing rAAV-based gene therapies.

## RESULTS

### Genome-wide screen of gRNA library to identify host restriction factors for rAAV5 transduction in 293-F cells

GPR108, localized to the TGN, facilitates AAV transporting from endosomes to the Golgi apparatus by interacting with AAV VP1u (35); however, AAV5 is unique in that it does not use GPR108 in this process. To identify additional host factors that restrict AAV5 intracellular trafficking, we conducted a genome-wide CRISPR/Cas9 screen in suspension 293-F cells using rAAV5, as outlined in **Figure 1A**. Flow cytometry was used to select the rAAV5-untransduced (mCherry−) cells, followed by two rounds of selection. Genomic DNA was then extracted for next-generation sequencing (NGS) and analyzed using MAGeCK software package (**Table S1**) (42). The NGS results revealed several host genes that were disrupted in the subset of rAAV5-untransduced cells were enriched in the first round of screening (Sort 1), compared to the unselected cells (Sort 0) (**Figure S1**, Sort 1-0), which was further enriched in second round of screening (Sort 2) (**Figure 1B**, Sort 2-0). These genes include the previously reported *KIAA0319L*, *TM9SF2*, and *RNF121*, which encode factors that restrict rAAV transduction (26,35–37), as well as a cluster of genes involved in glycan biogenesis, Golgi localization and ER penetration. While the known limiting factor, TM9SF2 is ranked at the top of the enriched genes of the mCherry− cells (resistant to AAV5 transduction), SLC35A1 is the runner-up when comparing Sort 2 vs Sort 0 (**Figure 1B**). It has not been identified or ranked high on previous screens (38,43).

**Figure 1.**
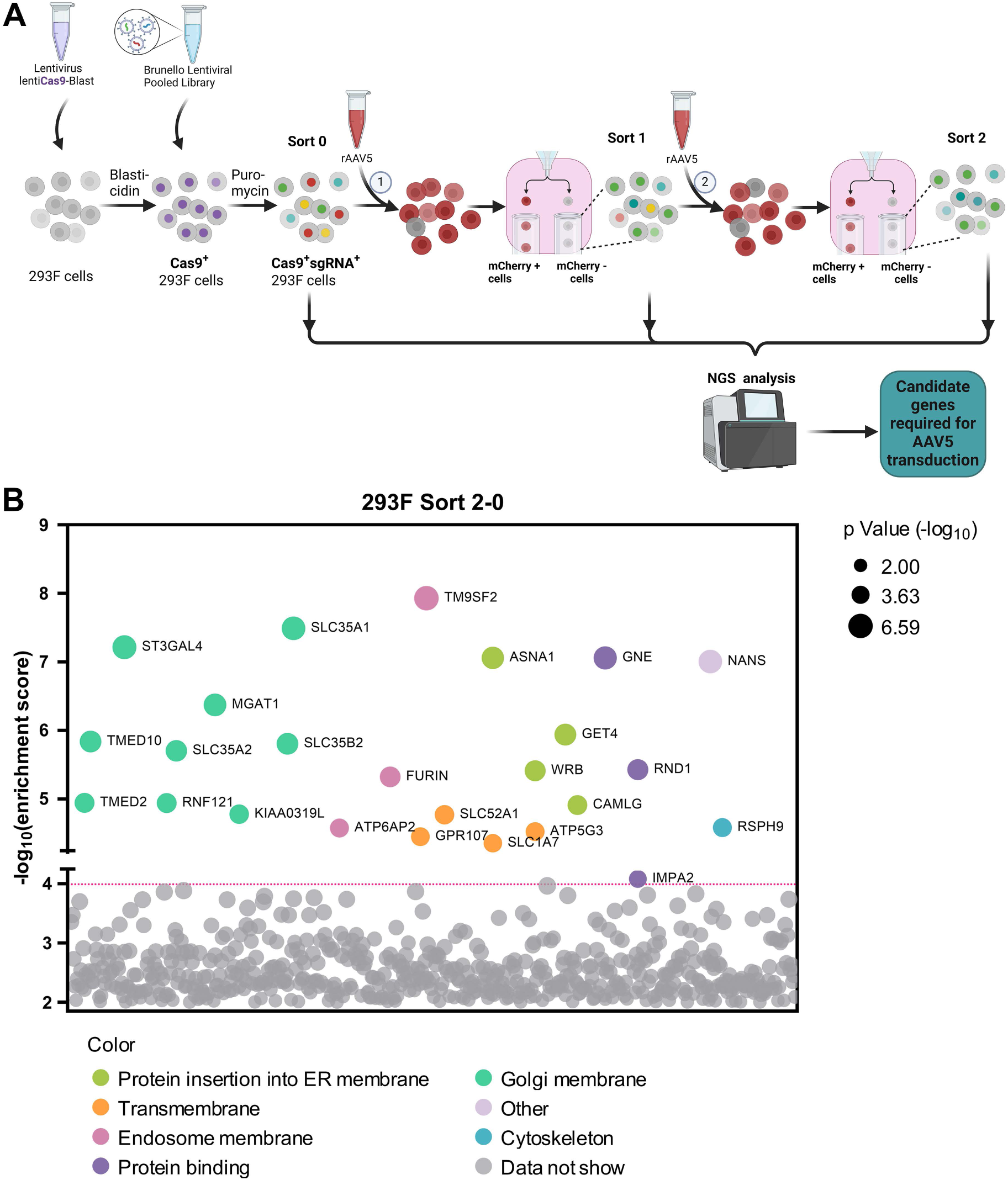
The Genome-wide CRISPR/Cas9 screen identifies host factors required for rAAV5 transduction. **(A) Diagram of genome-wide CRISPR/Cas9 gRNA library screen.** Suspension 293-F cells were transduced with a lentiviral vector carrying spCas9 and a blasticidin resistance gene followed by blasticidin selection. Blasticidin-resistant spCas9-expressing cells (1×10^8^) were then transduced with the Brunello lentiCRISPR gRNA lentiviral library and selected with puromycin to obtain Cas9/sgRNA-expressing 293-F cells. The selected cells were cultured and expanded to 2×10^8^. Among them, 1×10^8^ cells were harvested for genomic DNA (gDNA) extraction as the control (gDNA^Sort0^), while the other 1×10^8^ cells were transduced with mCherry-expressing rAAV5. Flow cytometry was performed at 3 days post-transduction (dpt), and the top 1% mCherry-negative (mCherry−) cells were collected and expanded to 2×10^8^ as the Sort 1 cells. We used 1×10^8^ cells from this population for gDNA extraction (gDNA^Sort1^), and another 1×10^8^ cells for the 2^nd^ round screening of rAAV5 transduction. The mCherry− cells from this round collected from cell sorting were expanded to 1×10^8^ for gDNA extraction (gDNA^Sort2^). The gDNA samples were subjected to NGS and bioinformatics analysis. **(B) Enrichment of genes from the 2^nd^ round screen of mCherry− cells.** NGS analyses were aimed at the sgRNA recognition sequences present in the mCherry− cell population, which identified the disrupted target genes at these sites. The x-axis represents genes targeted by the Brunello library, grouped by gene ontology analysis. The y-axis shows the enrichment score [-log_10_] of each gene based on MAGeCK analysis of the sgRNA reads in gDNA^Sort2^ vs. gDNA^Sort0^. Each circle represents a gene, with its size indicating the statistical significance [-log_10_] of enrichment when comparing gDNA^Sort2^ to gDNA^Sort0^. The color of each circle represents the function of the genes. Only genes with an enrichment score greater than 10^4^ are shown.

SLC35A1, localized to the Golgi apparatus, is a CMP-SIA transporter that plays a role in the biogenesis of SIA (39). We investigated how the *SLC35A*1 KO in HEK293 cells affected the transduction of rAAV2, rAAV5 and rAAV2.5T vectors. As a comparison, we included the KO cells of the known *TM9SF2* and *KIAA0319L* genes, as well as another gene, *TMED10*, encoding a component of COPII-coated vesicles that is required for efficient ER to Golgi transport (44). Target gene KO cell lines were made using an established CRISPR/Cas9 technique (38,45). Western blotting confirmed the absence of the corresponding proteins in cell lysates, validating the specific gene KOs (**Figure 2A**). Each cell line was then transduced by rAAV5, rAAV2, and rAAV2.5T, respectively, to compare the impacts of these gene KOs on transgene reporter expression, which was normalized to that from the scramble control cell line. The validation in rAAV5 transduction revealed that *SLC35A1* KO significantly reduced rAAV5 transduction to levels lower than that in *KIAA0319L* or *TM9SF2* KO cells (**Figure 2B**). *SLC35A1* KO also significantly reduced rAAV2 transduction more than the level seen in *KIAA0319L* KO cells (**Figure 2C**). We also tested rAAV2.5T, and the results showed that *SLC35A1* KO significantly reduced rAAV2.5T transduction to the levels observed in *KIAA0319L* and *TM9SF2* KO cells (**Figure 2D**). KO of *TMED10* moderately decreased transduction of rAAV2, 5, and 2.5T (**Figure 2B-D**).

**Figure 2.**
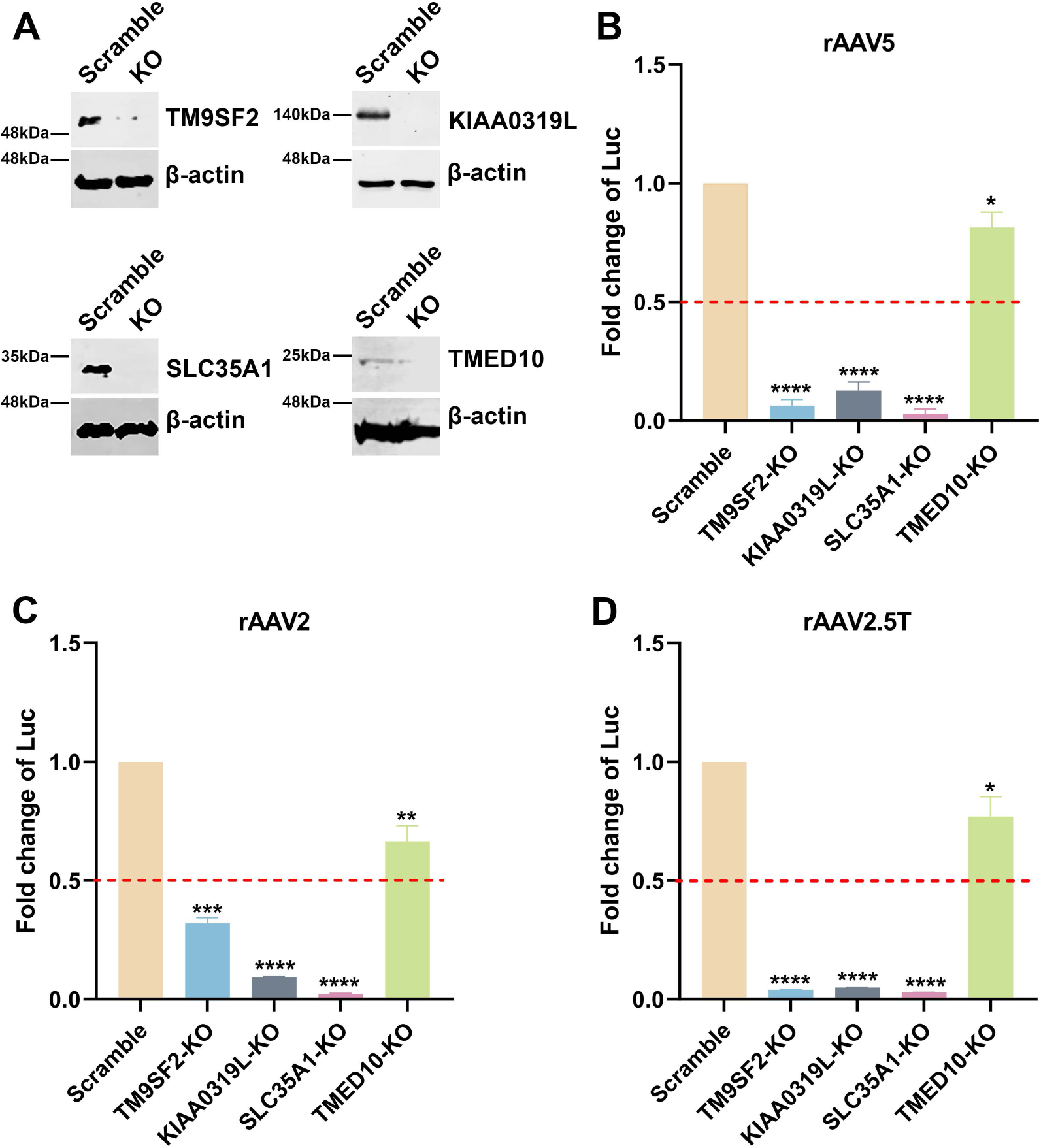
rAAV5, rAAV2, and rAAV2.5T transduction in *SLC35A1, TM9SF2, KIAA0319L,* and *TMED10* KO HEK293 cells. sgRNA-expressing lentiviral vectors were applied in HEK293 cells to generate *SLC35A1*, *TM9SF2*, and *KIAA0319L* KO cell lines. **(A) Western blotting.** Western blotting analysis shows the KO efficiency of scramble control and gene KO cells. β-actin was used as a loading control. **(B-D) Luciferase activities in gene KO HEK293 cells.** Scramble and gene KO cells were transduced with rAAV5 at an MOI of 20,000 DRP/cell (B), rAAV2 at an MOI of 2,000 DRP/cell (C) or rAAV2.5T at an MOI of 2,000 DRP/cell (D). At 3 dpt, the luciferase activities were measured. Data shown are the averaged luciferase activities relative to the Scramble cells from three replicates [mean plus standard deviation (SD)]. The red dashed line indicates 50% of the luciferase activity in Scramble HEK293 cells. P values were determined by using one-way ANOVA for the comparison of the fold changes in the KO cell groups and the Scramble cell control.

Taking these results together, we concluded that *SLC35A1* encodes an important factor that plays a direct or indirect role in the transduction of rAAV2, rAAV5, and rAAV2.5T as important as KIAA0319L in HEK293 cells. As *SLC35A1* involves the biogenesis of SIA and both rAAV5 and AAV2.5T utilizes α2,3-linked SIA as a primary attachment molecule, we further studied the mechanisms underlying the role of SLC35A1 in rAAV transduction.

### Effect of *SLC35A1* KO on SIA expression

To investigate the effect of SLC35A1 KO on the expression of various SIAs, we first used *Sambucus nigra* lectin (SNA) and *Maackia amurensis* lectin II (MAL II) to detect the expression of α2,6-and α2,3-linked SIAs, respectively (46,47). The results revealed that *SLC35A1* KO nearly abolished the overall expression of α2,6-linked SIA in cells, as shown by negative staining of SNA, but remained detectable expression of α2,3-linked SIA (47) (**Figure 3A&B**, *SLC35A1* KO). Cells treated with neuraminidase (NA) to remove all sialic acids were used as a negative control of lectin staining (**Figure 3A&B**, NA-treated). The expression of SIA on the cell surface was quantified by flow cytometry. The results showed that *SLC35A1* KO decreased α2,6-linked SIA by 90%, but only 64% in α2,3-linked SIA cell surface expression (**Figure 3C&D**).

**Figure 3.**
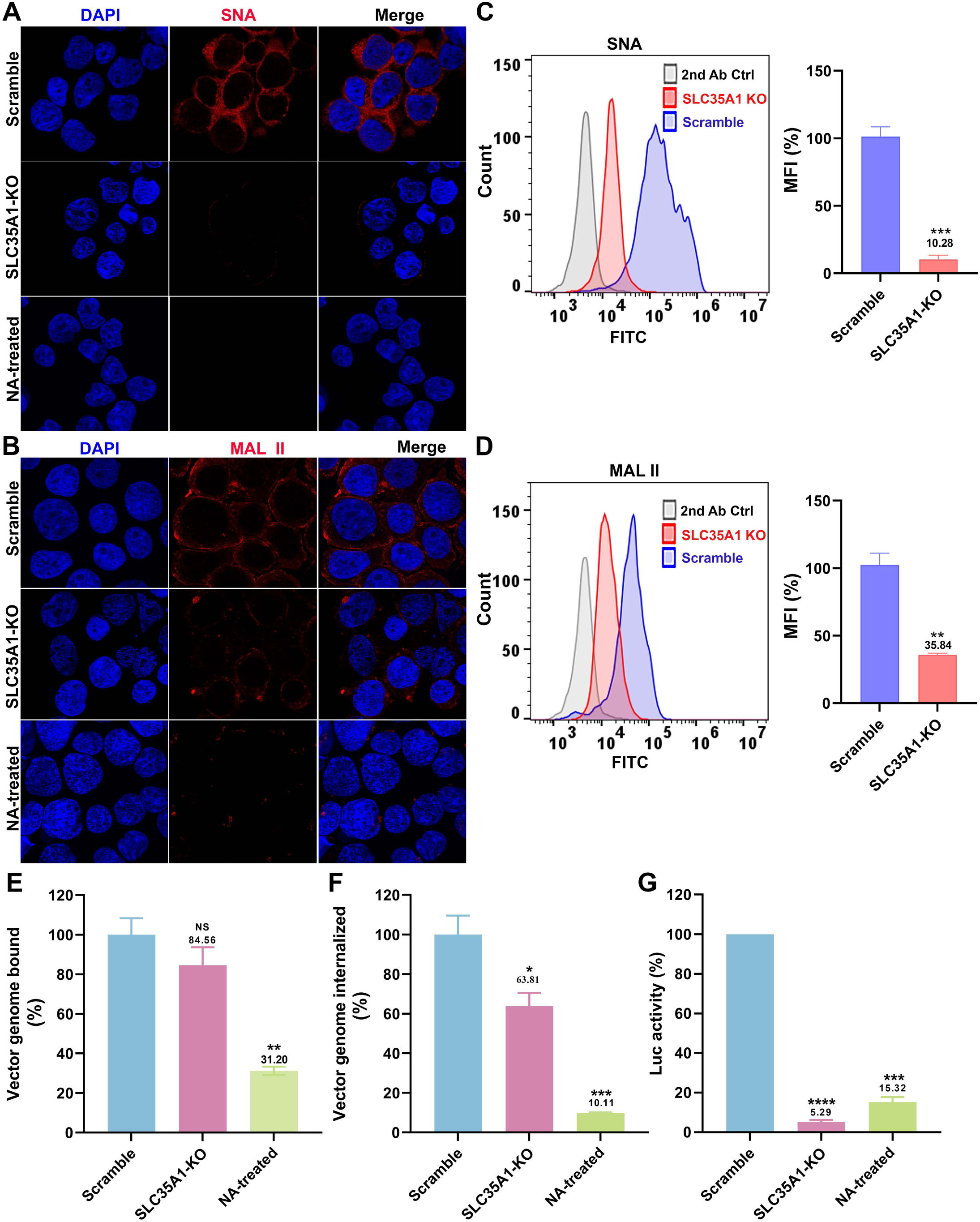
*SLC35A1* KO significantly decreases SIA expression in HEK293^SLC35A1-KO^ cells. **(A-F) Lectin staining.** Biotinylated *Sambucus Nigra* lectin (SNA) and *Maackia Amurensis* lectin II (MAL II) lectins were used to stain glycan expression in HEK293^SLC35A1-KO^ cells. NA-treated cells served as a positive control to show the removal of sialic acids. **(A&B) Confocal microscopy**. SNA (A) and MAL II (B) stained cells were incubated with DyLight 649-conjugated streptavidin for visualization at 100 × under a confocal microscope (Leica SP8 STED). **(C&D) Flow cytometry.** (C) SNA and (D) MAL II stained cells were incubated with FITC-conjugated streptavidin for flow cytometry. The histograms show the intensity of the FITC staining on the x-axis and the number of cells at each intensity level on the y-axis. The mean fluorescence intensity (MFI) values were calculated, normalized to the wild-type (WT) HEK293 cells as percentages (%), and are shown with a mean and SD from three replicates. P values were determined by using the Student’s *t*-test. **(E-G) rAAV5 vector transduction, binding, and internalization in HEK293 cells.** Relative percentages of vector binding (E), internalization (F), and transduction (G) to the Scramble cell group are calculated in rAAV-transduced SLC35A1-KO or NA-treated scramble HEK293 cells. The data shown were a mean and SD from three replicates. P values were determined by using one-way ANOVA for the comparison of the vector value in the KO or NA-treated cell group and the scramble cell group.

We next carried out rAAV binding, internalization and transduction assays in parallel in *SLC35A1* KO cells and NA-treated cells, with the transductions of the cells (Scramble) treated with a scramble-gRNA-expressing lentiviral vector as a control. The results showed that the *SLC35A1* KO did not significantly affect rAAV5 binding (**Figure 3E**), but significantly reduced rAAV5 entry by 36% (**Figure 3F**) and decreased transduction efficiency by 95% (**Figure 3G**). As a positive control, neuraminidase treatment decreased rAAV5 binding by 69%, internalization by 90%, and transduction by 85% (**Figure 3E-G**, NA-treated).

Overall, the KO of *SLCA35A1* did not significantly affect vector binding, likely because the KO only partially diminished the surface expression of α2,3-linked SIA, which rAAV5 used for cell attachment. However, the significant 36% decrease in vector entry, which may be due to the 15% less in vector binding, did not correlate with the 95% reduction in rAAV5 transduction. Thus, our findings suggest that while SLC35A1 is an essential host factor of rAAV5 transduction beyond the cell surface binding of the vector.

### *SLC35A1* KO reduces transduction of rAAV5 and rAAV2.5T in human airway epithelia

To further study the involvement of SLC35A1 in rAAV transduction, we investigated the transduction of rAAV5 and its airway-tropic variant rAAV2.5T in human airway epithelium culture at an air-liquid interface (HAE-ALI). HAE-ALI is a physiologically relevant model mimicking human airway epithelium and wildly used for studying the infections of respiratory viruses and airway gene transfer from viral vector transduction (48). To this end, we used CRISPR to disrupt the *SLC35A1* in CuFi-8 cells, an immortalized human airway cell line that retains the potential to differentiate into pseudostratified mucociliary epithelia when cultured at an ALI (49). rAAV5 and rAAV2.5T transductions were performed with the HAE-ALI cultures differentiated from *SLC35A1* KO CuFi-8 cells (HAE-ALI^SLC35A1-KO^) (**Figure S2A**). Before rAAV transduction assays, we performed several experiments to characterize these cultures. Western blotting confirmed no SLC35A1 expression (**Figure S2B**). Transepithelial electrical resistance (TEER) exceeding 1,800 Ω.cm^2^ indicated that neither *SLC35A1* KO nor NA-treatment impacted the epithelial integrity of the HAE-ALI cultures (**Figure S2C**), which was comparable to that of the *KIAA0319L* KO and Scramble HAE-ALI cultures. Lectin fluorescent staining microscopy showed that HAE-ALI^SLC35A1-KO^ cells barely expressed α2,6-linked SIA (**Figure 4A**), but some expression of α2,3-linked SIA (**Figure 4B**), consistent with the patterns observed in *SLC35A1* KO HEK293 cells (**Figure 3A&B**). NA-treated cultures served as positive controls for SIA removal. By cell surface flow cytometry, we confirmed a reduction of 87% in α2,6-linked SIA expression (**Figure 4C**) but only 28% in α2,3-linked SIA expression (**Figure 4D**).

**Figure 4.**
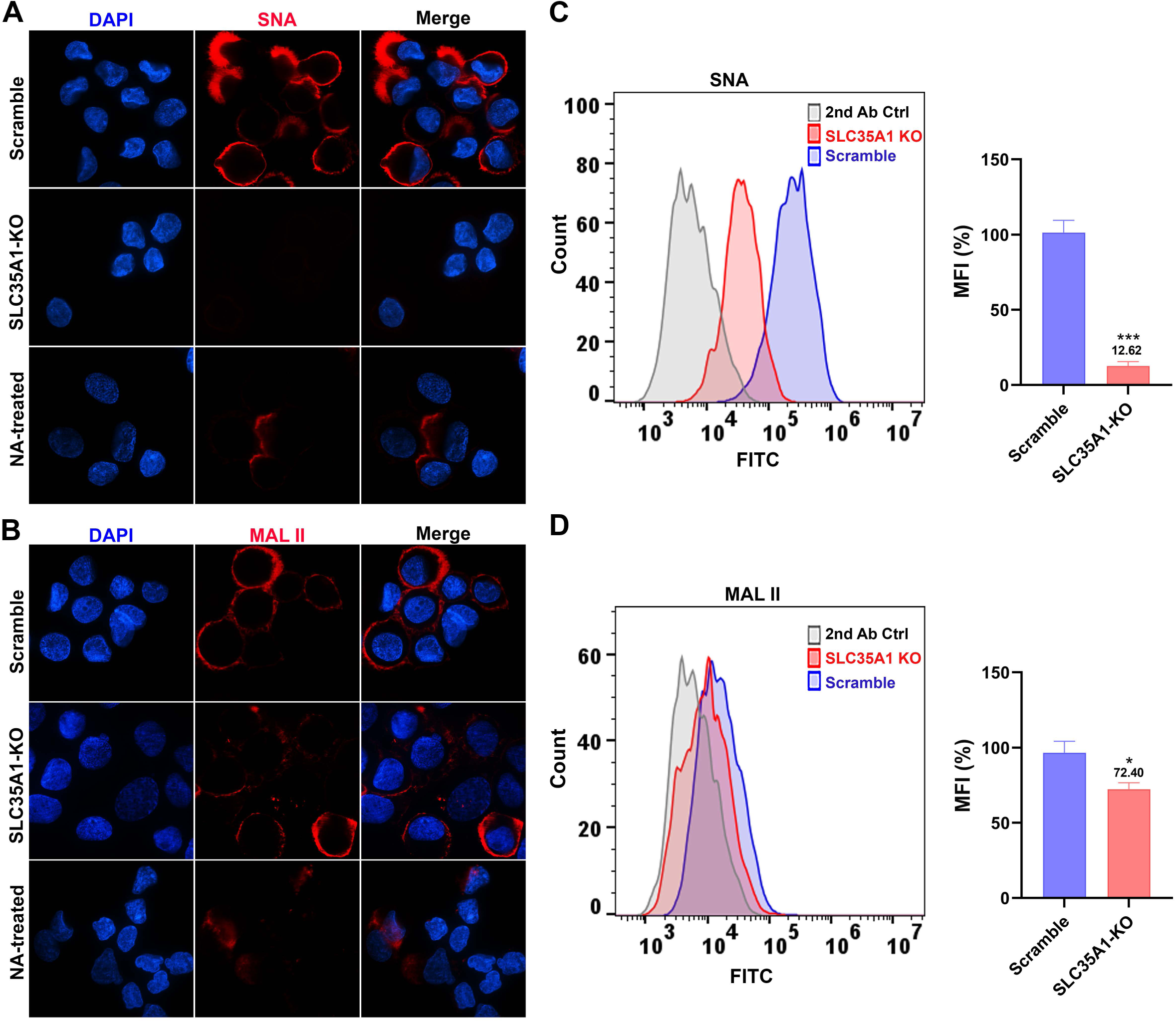
SIA expression in HAE-ALI cultures differentiated from *SLC35A1* KO cells. **(A&B) Confocal microscopy**. SNA (A) and MALII (B) lectins were used to stain glycan expression in HAE-ALI^SLC35A1-KO^ cultures. NA-treated cultures served as a positive control to show the removal of sialic acids. DyLight 649-conjugated streptavidin was used to visualize the staining under a confocal microscope at × 60 (CSU-W1 SoRa). **(C&D) Flow cytometry of lection-stained cells dissociated from ALI cultures.** (C) biotinylated SNA and (D) MAL II lectins were used to stain the cell surface, followed by FITC-conjugated streptavidin for detection. The histograms show the intensity of the FITC staining on the x-axis and the number of cells at each intensity level on the y-axis. The mean fluorescence intensity (MFI) values were calculated, normalized to the WT HEK293 cells. And the percentages (%) are shown with a mean and SD from three replicates. P values as indicated were determined by using the Student’s *t*-test.

We next examined the rAAV5 vector binding, internalization and transduction efficiency in HAE-ALI^SLC35A1-KO^ cultures. The KO of *SLC35A1* showed a 38% decrease in rAAV5 binding (**Figure 5A**) and a corresponding reduction in vector internalization by 26% (**Figure 5B**), but a drastic decrease in vector transduction by 98% (**Figure 5C**). In the transduction of rAAV2.5T, *SLC35A1* KO resulted in no significant decreases in vector binding and entry into the cells but a significant decrease in transduction by 76% (**Figure 5D-F**).

**Figure 5.**
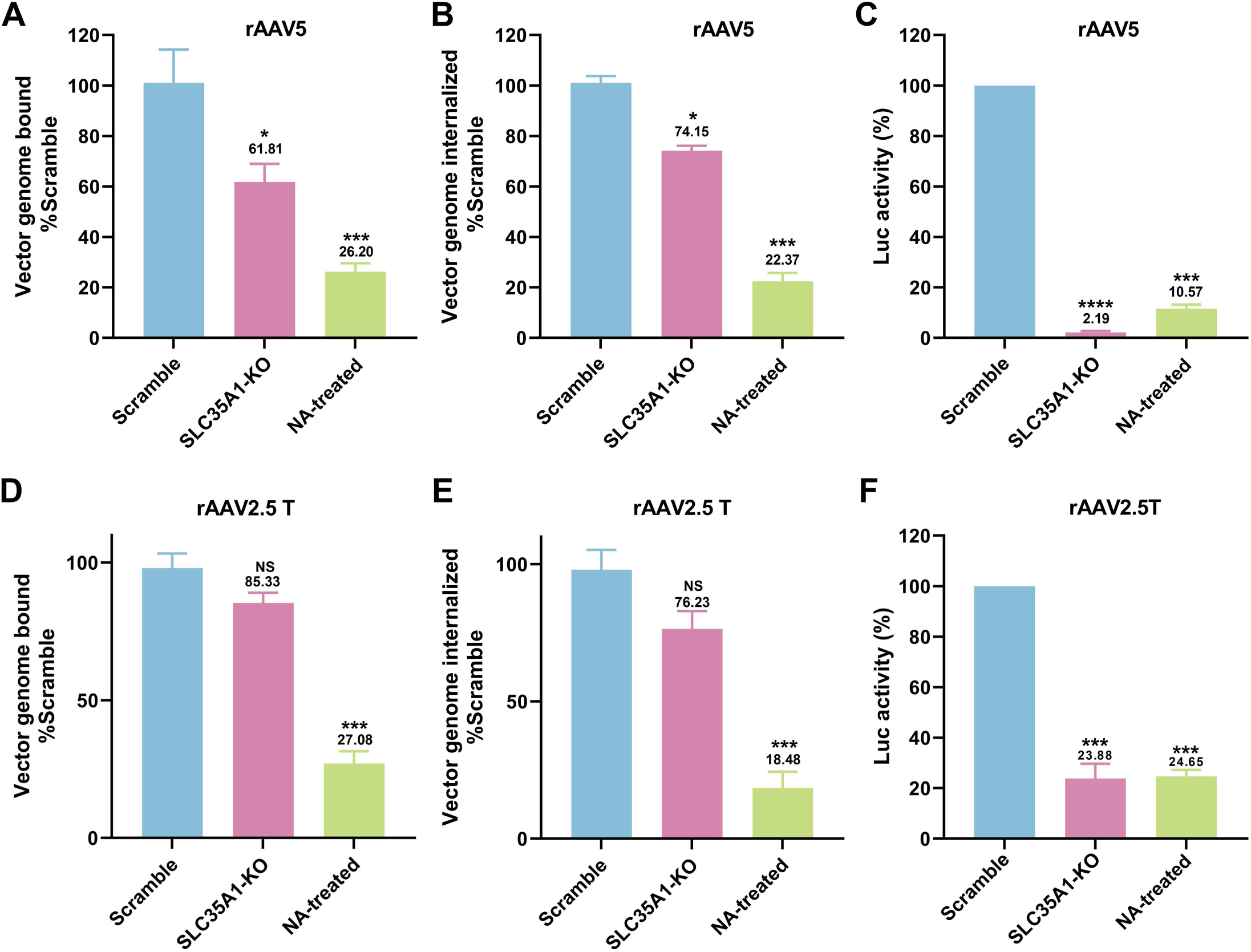
*SLC35A1* KO leads to a larger decrease in transduction efficiency than that of vector binding and entry of rAAV5 or rAAV2.5T in polarized HAE-ALI cultures. **(A-C) rAAV5 vector binding, internalization, transduction in HAE-ALI^SLC35A1-KO^ cultures.** HAE-ALI Scramble, *SLC35A1* KO or NA-treated scramble cultures were apically transduced with rAAV5 at an MOI of 20,000 DRP/cell. At 2 hpt, vector binding and internalization assays were carried out, and at 5 dpt, the luciferase activities were measured. Relative percentage of binding (A), internalization (B) or transduction efficiency (C) of the transduced KO and NA-treated cultures to the Scramble cell group are shown. **(D-F) rAAV2.5T vector transduction, binding, and internalization in HAE-ALI^SLC35A1-KO^ cultures.** The ALI cultures as indicated were transduced with rAAV2.5T at an MOI of 20,000 DRP/cell. Relative percentages of vector binding (D), internalization (E), and transduction efficiency (F) of the transduced KO and NA-treated cell cultures to the Scramble cell group are shown. All repeated data are shown with a mean and SD of at least three replicates. P values as indicated were determined by using one-way ANOVA for the comparison of the vector value in the KO or NA-treated cell group with the Scramble cell group.

Collectively, our data demonstrated in polarized human airway epithelium, SLCA35A1 is crucial for rAAV5 transduction across all stages, including binding and post-entry processing. In contrast, for rAAV2.5T transduction, SLCA35A1 plays a significant role primarily after vector internalization.

### *SLC35A1* KO significantly diminishes the transduction of rAAV1-8, rAAV12 and rAAV13, but increases the transduction of rAAV9 and rAAV11 in HEK293 cells

For a potentially broad role of SLC35A1 in rAAV transductions, we investigated its function in the transduction of various serotypes of rAAV in HEK293 cells by comparing *SLC35A1* KO and Scramble-treated HEK293 cells. The results showed that *SLC35A1* KO decreased the transduction of rAAV1-8, rAAV12 and rAAV13, but increased the transduction of rAAV9 and rAAV11 (50) (**Figure 6A**). It was reported that low or no SIA expression resulted in an elevated level of galactose-associated glycan presentation on the cell surface (50). The increased transduction of rAAV9 was likely due to the higher expression of galactose in *SLC35A1* KO cells that was confirmed by *Erythrina cristagalli* lectin (ECL) staining (**Figure 6B&C**).

**Figure 6.**
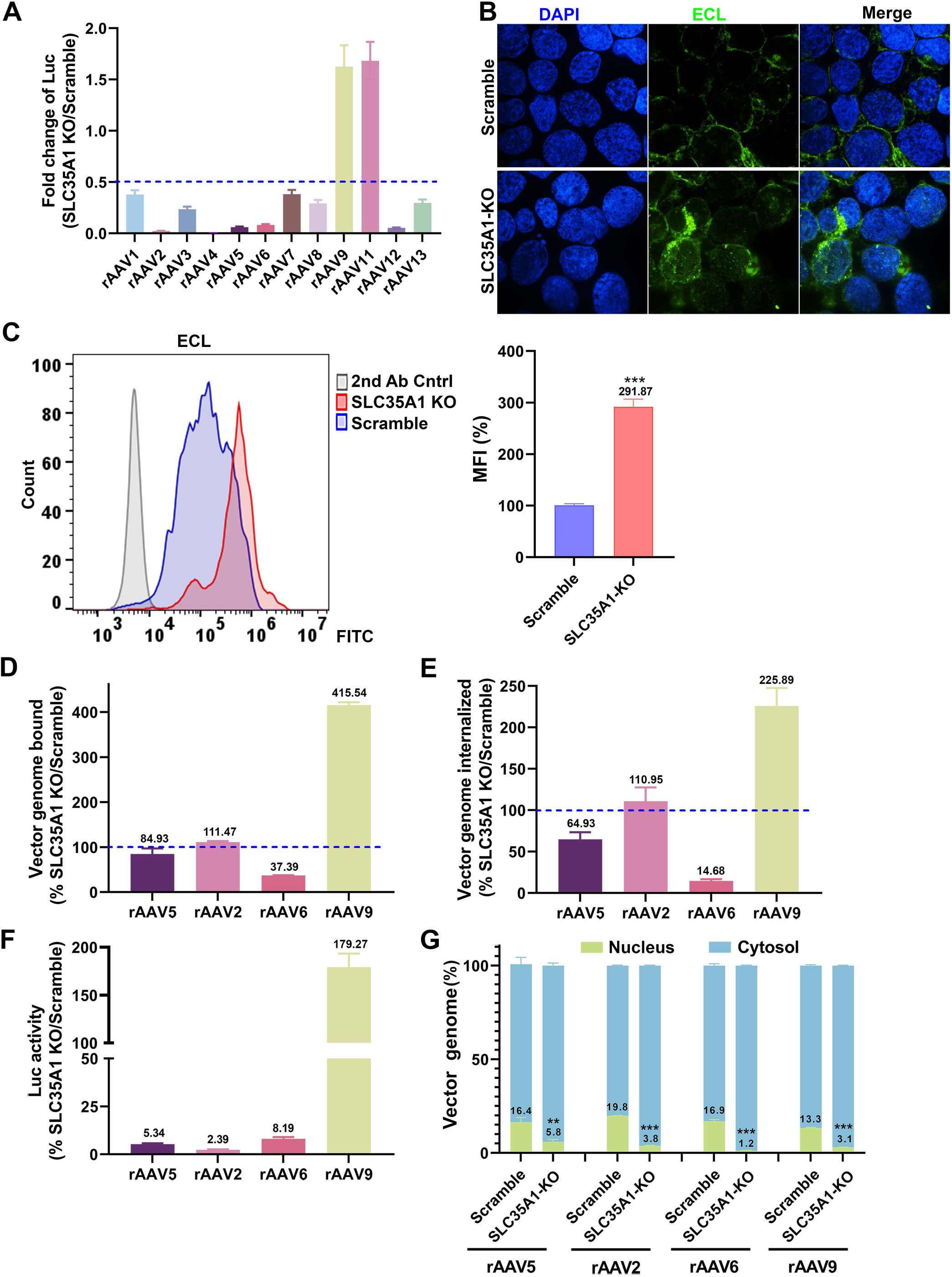
*SLC35A1* KO significantly decreases the transduction efficiency of rAAV1-8, 12 and 13, but increases the transduction efficiency of rAAV9 and rAAV11, and causes a significant decrease in the nuclear import of rAAV. **(A) Luciferase activities.** HEK293^Scramble^ and HEK293^SLC35A1-KO^ cells were respectively transduced with various serotypes of rAAV vectors as indicated at an MOI of 20,000 DRP/cell. At 3 dpt, the luciferase activities were measured and normalized to Scramble cells (set as 1). The fold changes of luciferase activities in SLC35A1-KO vs Scramble are shown with means and an SD from at least three replicates. **(B&C) Lectin staining.** HEK293^Scramble^ and HEK293^SLC35A1-KO^ cells were respectively stained with *Erythrina cristagalli* lectin (ECL) for analyses by confocal microscopy (B) and by flow cytometry (C). The mean fluorescence intensity (MFI) values were calculated and normalized to the Scramble cells as percentages (%), which are shown with a mean and SD from three replicates, and were analyzed by the Student’s *t*-test. **(D-F) rAAV binding, internalization, transduction.** HEK293^Scramble^ and HEK293^SLC35A1-KO^ cells were respectively transduced with four selected representative vectors, rAAV5, rAAV2, rAAV6, and rAAV9, in parallel. Vector binding (D), Internalization (E), and transduction (F) are assessed and relative fold changes in SLC35A1-KO vs Scramble as percentages (%) are shown with a mean and SD from at least three replicates. **(G) Nuclear import assays**. HEK293^Scramble^ and HEK293^SLC35A1-KO^ cells were respectively transduced with four selected representative vectors, rAAV5, rAAV2, rAAV6, and rAAV9, in parallel. At 12 hpt, the cytoplasm and nucleus were fractionated, and the percentage of vector genome copies in the cytoplasm and nucleus fractions were quantified. The data shown are means with an SD from at least three replicates. P value was determined by using the Student *t*-test for the comparison of the vector genome copies in the nucleus between the KO cell group and the Scramble cell group.

### *SLC35A1* KO significantly reduces binding and entry of rAAV6 but not rAAV2, and reduces nuclear import of rAAV2, rAAV5, rAAV6, and rAAV9 in HEK293 cells

We then examined the binding and internalization of the representative rAAV vectors, rAAV2, rAAV6, and rAAV9, along with rAAV5 as a comparison. Among these AAVs, AAV5 primarily utilizes α2,3-linked SIA (23,24), AAV2 uses heparan sulfate (16), AAV6 mainly binds to α2,6-linked SIA (21), and AAV9 uses N-linked galactose (20) for attachment. The data indicated that *SLC35A1* KO drastically reduced the binding and internalization of rAAV6 (**Figure 6D&E**), which correlated with the decreased transduction (**Figure 6F**), supporting the primary receptor role of α2,6-linked SIA in rAAV6 transduction (21), as *SLC35A1* KO removed α2,6-linked SIA by 90% on the cell surface (**Figure 3A&C**). On the other hand, *SLC35A1* KO increased both the binding and internalization of rAAV9, correlating with increased transduction (**Figure 6D-F**), due to the increased expression of galactose (**Figure 6B&C**). Importantly, *SLC35A1* KO had no effects on the binding and internalization of rAAV2, while the transduction efficiency was significantly reduced (by 98%) (**Figure 6D-F**), supporting our hypothesis that SLC35A1 primarily acts in post-entry processing of rAAV.

To investigate the effects of SLC35A1 on the intracellular trafficking of the internalized vectors, we assessed the viral genome distributions in the cytoplasm and the nucleus by cell fractionation. The results showed that *SLC35A1* KO significantly reduced the nuclear import of rAAV5 and rAAV2 by ∼3-and 5-fold, respectively, and the nuclear import of rAAV6 by 12-fold (**Figure 6G**). Importantly, although the rAAV9 transduction was notably increased in *SLC35A1* KO cells (**Figure 6A**), the KO significantly decreased the efficiency in nuclear import of rAAV9, as shown by the 4-fold lower level of the vector in the nucleus of the *SLC35A1* KO cells (**Figure 6G**).

We then examined the localization of SLC35A1 in rAAV-transduced cells by immunofluorescent assays and confocal microscopy. At an early time point of 8 hours post-transduction (hpt), SLC35A1 was primarily localized to the TGN, visualized by co-localization with TGN46 (**Figure 7A**). When looking at the details of the rAAV5 capsid in the nuclei, the capsids were rarely found in the nuclei of *SLC35A1* KO cells, compared to the rich staining of AAV5 capsids in the nuclei of scrambled control cells (**Figure 7A**), supporting that *SLC35A1* KO impeded vector nucleus import (**Figure 6G**). Furthermore, we observed that SLC35A1 colocalized with the rAAV2.5T capsid, and the capsids were rarely found in the nuclei of *SLC35A1* KO HEK293 cells, compared to scrambled controls (**Figure 7B**), which was further confirmed in the cells of HAE-ALI^SLC35A1-KO^ cultures (**Figure 7C**).

**Figure 7.**
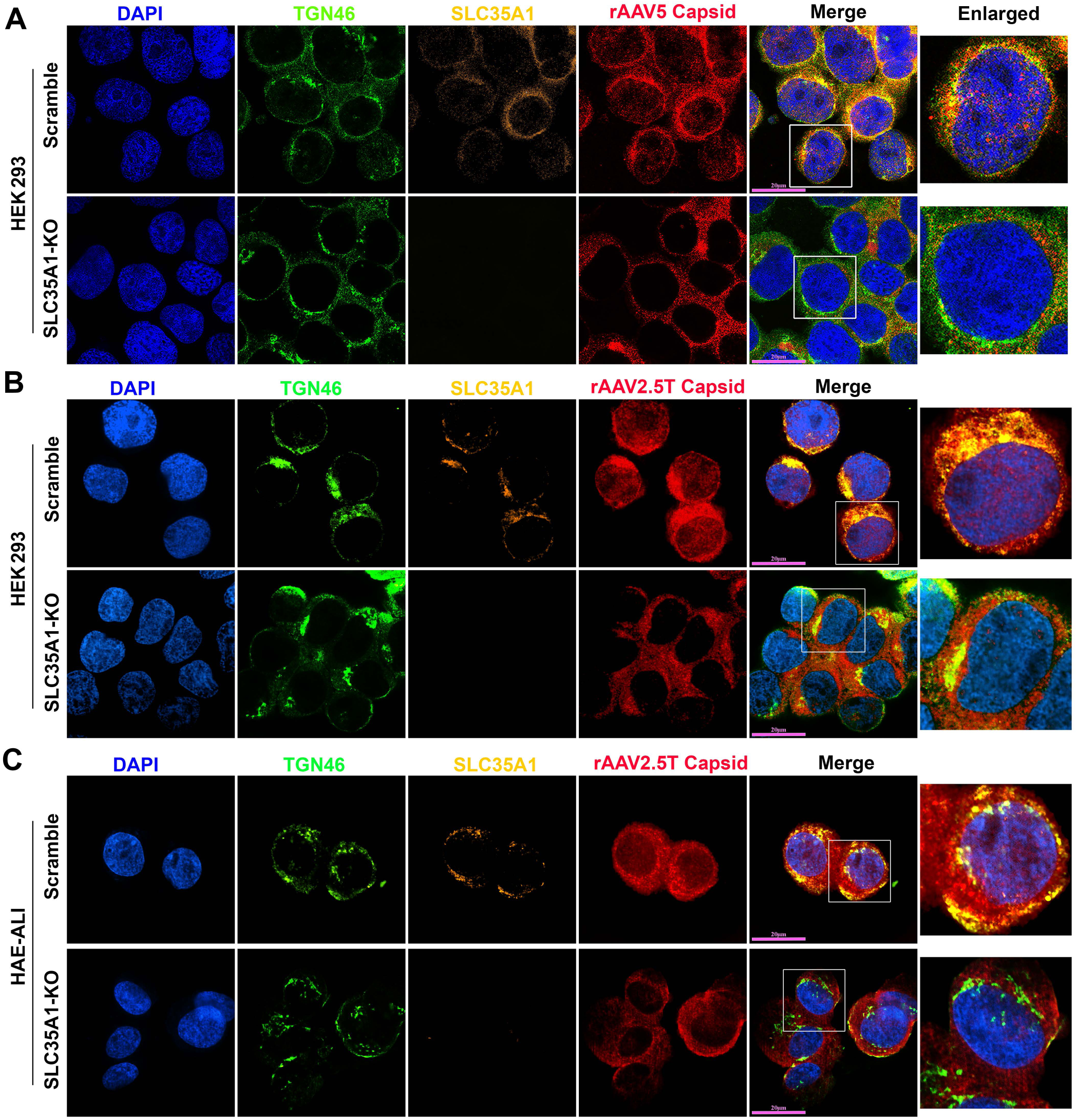
SLC35A1 and AAV capsid are colocalized with TGN46. **(A&B) HEK293 cells.** SLC35A1-KO or Scramble HEK293 cells were transduced with (A) rAAV5 at MOI of 20,000 or (B) rAAV2.5T at MOI of 2,000. At 8 hpt, the cells were fixed and permeabilized, followed by immunostaining with the first antibody against indicated protein and fluorescence-conjugated secondary antibodies. **(C) HAE-ALI cultures.** The HAE-ALI cultures differentiated from SLC35A1-KO or Scramble CuFi-8 cells were transduced with rAAV2.5T at MOI of 20,000. At 3 dpt, the cells were fixed and permeabilized, followed by immunostaining with the first antibody against indicated protein and fluorescence-conjugated secondary antibodies. The stained cells were imaged under a confocal microscope (CSU-W1 SoRa, Nikon) at 60× with 4×SoRa magnitude (scale bar = 20 μm).

Collectively, our data demonstrated that SLC35A1 plays an important role in the nuclear import of rAAV5, 2, 6 and 9 in HEK293 cells, as well as the nuclear import of rAAV2.5T in HAE-ALI. Considering the TGN localization of SLC35A1, the results suggest that SLC35A1 plays an important role in the trafficking of multi-serotype AAVs to the TGN after internalization. Thus, our results categorized the important role of SLC35A1 in the transduction of rAAV in four groups: 1) rAAV2-type, in post-entry trafficking; 2) rAAV5-type, minor in vector binding to SIA and major in post-entry trafficking; 3) rAAV6-type, in both vector binding to SIA and post-entry trafficking; 4) AAV9, in vector binding (to galactose-associated glycan) and post-entry trafficking.

### The C-terminal tail of SLC35A1 is essential for rAAV nuclear import

Studies have shown that a mutation of T128 in SLC35A1, which is in the central sugar pocket, remained correct localization to the Golgi but became deficient in SIA biogenesis (51). The C-terminal cytoplasmic tail (C-Tail; 20 aa) is required for SLC35A1 to exit from the ER and localize to the Golgi, and the C-tail deletion mutant (ΔC Tail) does not localize to the Golgi (52). To differentiate the functions of SLA35A1 between SIA biogenesis and rAAV transduction, we used the T128A and ΔC Tail mutants to assess the functional complementation for SIA expression and rAAV transduction in *SLC35A1* KO cells.

We found that the ΔC Tail mutant was unable to restore the decreased transduction and nuclear import of rAAV5 in HEK293 cells caused by *SLC35A1* KO, whereas the T128A mutant fully restored the transduction accompanied by a restoration of nuclear import, in line with the Scramble control (**Figure 8A&B**). Meanwhile, we monitored SIA expression of the *SLC35A1* KO cells complemented with wild-type (wt)SLC35A1, T128A and ΔC Tail mutants, using flow cytometry for cell surface and immunofluorescent assay for intracellular expression, respectively. The ΔC Tail mutant restored 65% and 73% of the expression of α2,3-and α2,6-linked SIAs, respectively, on the surface of HEK293^SLC35A1 KO^ cells, but the T128A mutant poorly restored the SIA expression (**Figure 8C-F**). These results were consistent with the SIA expression in these cells observed in the immunofluorescent assays. The ΔC Tail mutant remained expression of SIAs in the cells, whereas the T128A barely expressed any SIAs (**Figure S3**).

**Figure 8.**
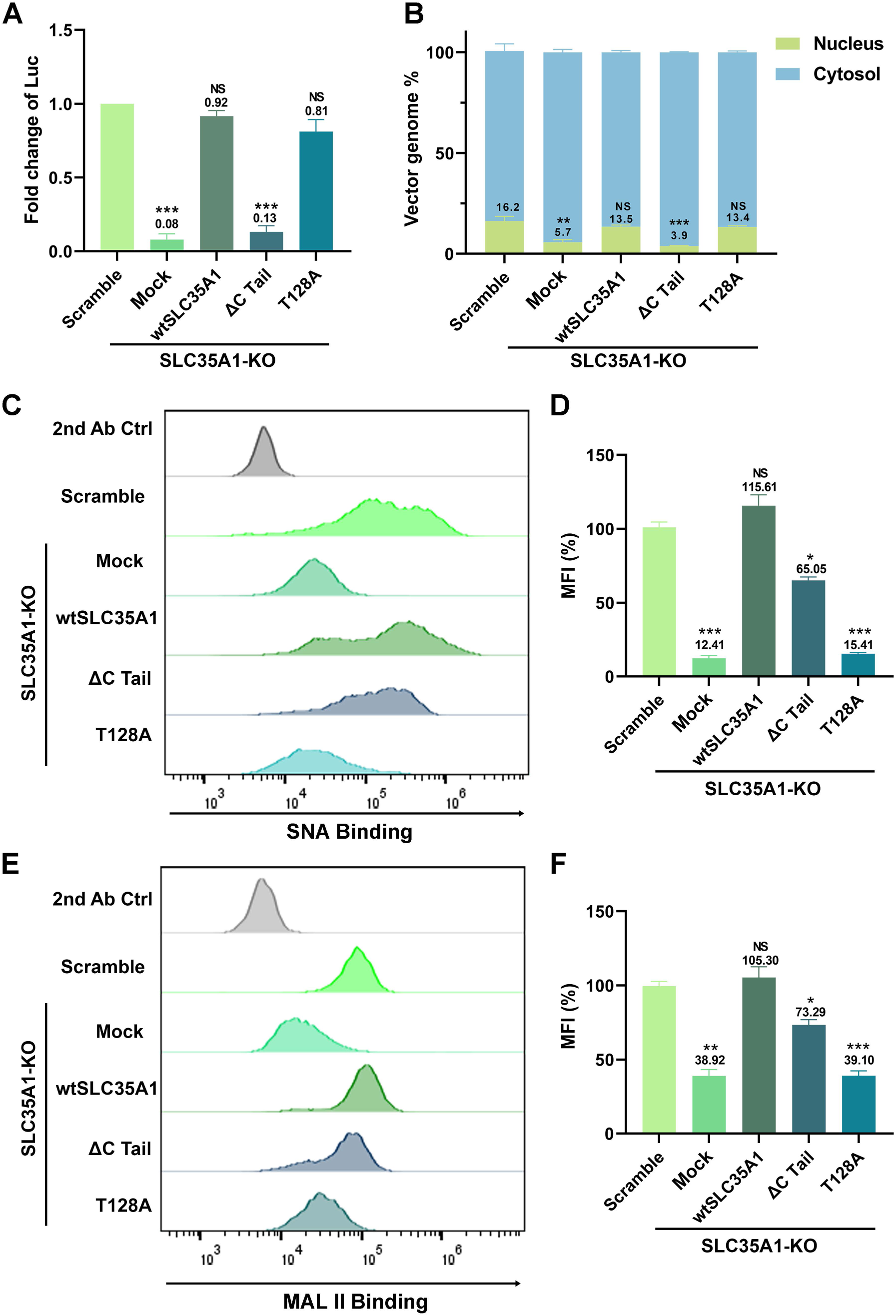
Expression of SLC35A1 wild-type (WT) and T128A mutant restores rAAV5 transduction and nuclear import, but not the ΔC Tail mutant in HEK293^SLC35A1^ cells. HEK293^SLC35A1-KO^ cells were transduced with lentiviral vector that expressed SLC35A1 WT, T128A and ΔC Tail, as indicated, or untransduced (Mock), followed by selection of blasticidin (at 10 µg/ml) for a week. The blasticidin-resistant cells were transduced with rAAV5 at an MOI of 20,000. HEK293^Scramble^ cells were used as a control. **(A) rAAV transduction efficiency.** At 3 dpt, luciferase activities were measured and normalized to the Scramble (set as 1.0). Data shown are means with an SD from three replicates. P values were determined by using one-way ANOVA for the comparison of the fold changes in the SLC35A1 KO cell groups and the Scramble cell control. **(B) rAAV genome distribution.** After 12 hpt, nuclear and the cytoplasmic fractions of the rAAV5-transduced were fractionationed, and the vector genomes in each fraction were quantified by qPCR. The percentage of viral genome in each fraction shown are means with an SD of three replicates. P values were determined by using one-way ANOVA for the comparison of the vector genome copies in the nucleus between the SLC35A1 KO cell groups and the Scramble cell control. **(C-F) Flow cytometry of lectin staining**. The cells were stained with biotinylated SNA (C&D) or MALII (E&F) lectin and FITC-conjugated streptavidin, followed by flow cytometry. The mean fluorescence intensity (MFI) values were calculated, normalized to the WT HEK293 cells as percentages (%), and shown as means with an SD from at least three replicates. P values were determined by using one-way ANOVA for the comparison of the fold changes in the SLC35A1 KO cell groups and the Scramble cell control.

Taken together, these results suggest that the C-terminal tail of SLC35A1 is essential for rAAV transduction, which is particularly important for vector nuclear import, where the cell surface expression of both α2,3-and α2,6-linked SIAs appeared not drastically influenced by the deletion of the C tail. Importantly, as the ΔC Tail mutant retains SIA expression and the T128A mutant is defective in SIA synthesis, the function of SLC35A1 as a CMP-SIA transporter can be differentiated from its function in the nuclear import of rAAV.

## DISCUSSION

We conducted a comprehensive genome-wide CRISPR/Cas9 screen to identify essential host factors for rAAV5 transduction in suspension 293-F cells. Our screen successfully identified several known AAV transduction restriction factor genes, such as *KIAA0319L* and *RNF121* (26,35–37), but notably did not identify GPR108 (35), validating the effectiveness of our screening approach. Significantly, our screen revealed a cluster of genes involved in ER penetration, including ER-anchoring factor gene *ASNA1* and an ER membrane gene *WRB* (53), as well as a cluster of genes related to glycan biogenesis, including *TM9SF2* (heparan sulfate synthesis), *SLC35A1* (CMP-SIA transporter), *ST3GAL4* (the main α2,3-sialyltransferase acting on N-glycans), *GNE* (biosynthesis of N-acetylneuraminic acid (NeuAc), a precursor of SIA), and *MANS* (a glycosyltransferase). Notably, except for *TM9SF2*, these glycan biogenesis genes have not been identified in previous screens. The discovery of these novel genes underscores the crucial role of ER penetration and glycosylation in AAV5 transduction. While further investigation is warranted to fully understand the roles these genes play in rAAV transduction, our study focused on SLC35A1. Surprisingly, we discovered that SLC35A1 is essential for the transduction of multiple rAAV serotypes except for rAAV9 and rAAV11. Beyond its role in SIA biogenesis, which is important for rAAVs that use SIA-based glycans as attachment receptors, SLA35A1 also facilitates vector nuclear import after vector internalization. Given its localization in the TGN, SLA35A1 likely aids in the trafficking of rAAV vectors through the TGN, thereby facilitating their nuclear import.

SLC35A1, a CMP-SIA transporter localized to the Golgi apparatus, plays a critical role in SIA-based glycan biogenesis (52). Our data demonstrated that knockout of *SLC35A1* in HEK293 cells significantly (>90%) reduced the presence of α2,6-linked SIA on the cell surface, while retaining a moderate (∼36%) expression of α2,3-linked SIA, as previously reported (54,55). This reduction in α2,3-linked SIA expression correlated with a ∼20-30% decrease in the binding and entry of AAV5, as AAV5 uses α2-3 N-linked SIA as a primary attachment receptor (24,25). However, the substantial reduction in rAAV5 transduction (∼95%) observed in *SLC35A1* KO cells was not proportional to the number of internalized vectors, suggesting that SLC35A1 influences the post-entry steps during rAAV5 transduction rather than merely facilitating its initial binding. This hypothesis is further supported by the experiments in polarized human airway epithelial ALI cultures, where *SLC35A*1 KO resulted in a ∼98% decrease in rAAV5 transduction but was associated with only a ∼25% reduction in vector internalization. Furthermore, in the scenario of rAAV2.5T, SLA35A1 primarily affects the transduction without significantly altering vector binding and entry.

AAV2 uses HSPG as an attachment receptor (16), and therefore the KO of SLAC35A1 did not affect both binding and entry. However, as AAV6 uses both α2-3 and α2–6 N-linked SIAs for attachment (21,22), KO of *SLC35A1* decreased both binding and entry of AAV6. Notably, KO of *SLC35A1* significantly enhanced the transduction of AAV9, which is correlated to the increase in vector binding and entry. The increase in AAV9 transduction upon *SLC35A1* knockout is due to the higher exposure of galactose residues, compensating for the loss of 2,6-linked sialic acid (50). Overall, the KO of *SLC35A1* significantly decreases the transduction efficiency of various other AAV serotypes, including AAV1-8, 12, and 13, while increasing the transduction of AAV9 and AAV11. This broad impact highlights not only the central role of SLC35A1-mediated sialylation in AAV binding but also an important role post-entry.

The cell fractionation and immunofluorescence assays provided deeper mechanistic insights into the post-entry role of SLC35A1 in rAAV transduction. We observed a significant reduction in the nuclear import of AAV2, 5, 6, and 9 in SLC35A1 KO cells, suggesting that SLC35A1 is additionally involved in the intracellular trafficking of AAV to the nucleus. The co-localization of SLC35A1 with AAV capsids in the Golgi apparatus marked by TGN46 supports the hypothesis that SLC35A1 facilitates the transport of AAV through the TGN, which assists in its nuclear entry (**Figure 9**). Thus, SLC35A1, which transports CMP-SIA from the cytosol into the Golgi apparatus lumen (39), contributes to the Golgi apparatus-loading of AAV vectors of both GPR108-dependent (AAV2-type) and independent types (AAV5-type) (**Figure 9**). The role of SLC35A1 in rAAV nuclear import transduction is also evident in polarized human airway epithelia. While *SLC35A1 KO* only decreased AAV5 internalization by 25%, it led to a dramatic 98% reduction in rAAV5 transduction. This disproportional effect between vector entry and transgene expression was also observed with rAAV2.5T. In HAE-ALI^SLC35A1-KO^, the internalization of rAAV2.5T decreased by 25%, but vector transduction dropped by 75%, accompanying fewer AAV2.5T capsids detected in the nucleus (**Figure 7C**). Notably, AAV2.5T is a variant of AAV5 with enhanced airway tropism, it transduced HAE-ALI much more efficiently than the parent rAAV5 (9).

**Figure 9.**
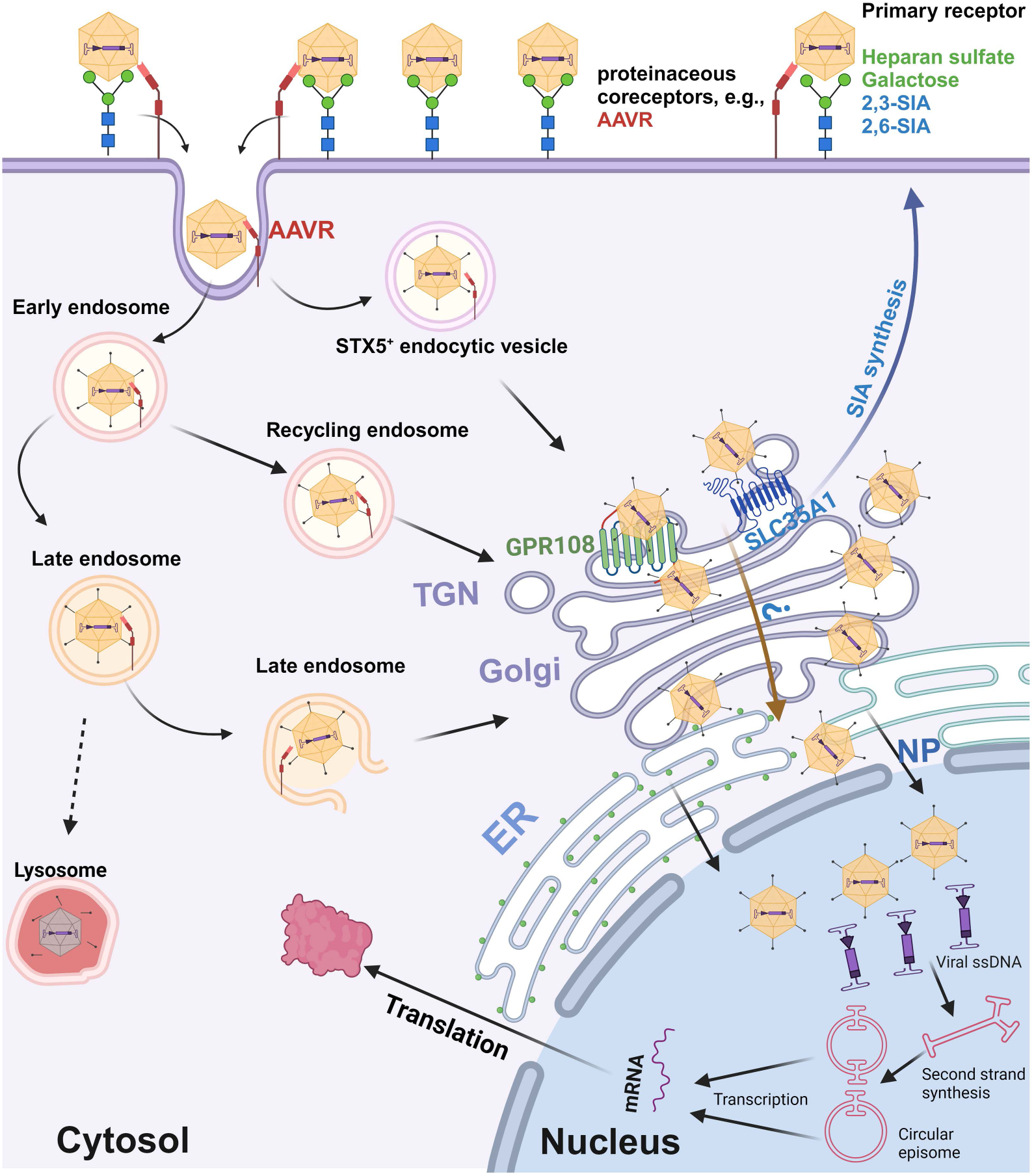
A model of SLC35A1 function in rAAV transduction. AAV cell entry is initiated by interacting with specific glycan on the cell surface (primary attachment receptor) (15,16,18,19,21,23) and a proteinaceous receptor, e.g., AAVR (KIAA0319L) (26,28). Several intracellular trafficking pathways have been proposed based on AAV2 studies. Post endocytosis or internalization, AAV traffics through Rab7^+^ late endosomes, Rab11^+^ recycling endosomes (64), and the STX5^+^ endocytic vesicle (65), to the TGN (66–68), where GPR108 localized (35), as well as SLC35A1. We hypothesize that SLC35A1, which transports CMP-SIA from the cytosol into the Golgi apparatus lumen (41), mediates AAV transport from the cytosol into lumen of the Golgi apparatus in a GPR108-dependent (AAV2-type) or independent (AAV5-type) manner AAVs, which likely facilitates vector nuclear import. Then, AAV traffics through the Golgi apparatus to the nuclear membrane and enters the nucleus through the nuclear pore (NP), or routes to a nonproductive pathway, e.g., proteasome, for degradation (not shown). In the nucleus, AAV releases the ssDNA genome, which is converted to dsDNA intermediates (1). The dsDNA further undergoes intra/intermolecular recombination of the inverted terminal repeats (ITRs) to form either linear or circular episomes that are transcribed to produce mRNA. Created in BioRender.

In the functional complementation assays in SLC35A1 KO cells, we demonstrated that the wtSLC35A1 fully restored both SIA expression and rAAV transduction. Interestingly, the expression of a T128A mutant fully compensated for the loss of rAAV transduction and facilitated AAV nuclear import in SLC35A1 KO cells, despite its inability to complement SIA biosynthesis, consistent with the previous report that T128 is important for the CMP-SIA transporter activity of SLC35A1 (51). These results support the role of SLA35A1 as a critical factor in AAV intracellular trafficking, independent of its role in SIA biosynthesis; therefore, the function of SLC35A1 in AAV intracellular trafficking is unlikely mediated by the changes in the properties of cellular glycans. Remarkably, expression of the ΔC Tail mutant, which lacks the C-terminal cytoplasmic tail, failed to restore rAAV5 transduction in SLC35A1 KO cells. The C-terminal cytoplasmic tail, consisting of only 20 aa, is required for SLC35A1 to exit the ER and localize to Golgi (52). This outcome highlights the importance of the Golgi localization in rAAV trafficking, as the transit of rAAV through the TGN is an essential process for its subsequential nuclear import. Nevertheless, our results still could not rule out the possibility that SLC35A1 may indirectly involve in AAV vector intracellular trafficking/nuclear import by affecting other host proteins that directly interact with AAV, which remains further investigation (**Figure 9**).

Overall, the identification of SLC35A1 and its universal role in AAV transduction has significant implications for advancing AAV-based gene therapies. Understanding the specific host factors required for AAV5 infection and other serotype vectors can inform the design of more effective vectors and enhance their transduction efficiency in target tissues. Additionally, manipulating glycosylation pathways, particularly those involving SLC35A1, could enhance the delivery and expression of therapeutic genes. Future studies should explore the detailed mechanisms by which SLC35A1 and other identified factors (yet to be investigated in this report), i.e., the ER-penetration factors, facilitate AAV trafficking and nuclear entry. This could lead to the development of optimized AAV vectors for clinical applications. By leveraging the insights from this study, we can refine the vector design and delivery strategy for gene therapies, ultimately advancing the treatment of genetic diseases.

## MATERIALS AND METHODS

### Cells and cell culture

#### HEK293 cells

HEK293FT (#R70007,ThermoFisher Scientific, Waltham, MA) and HEK293 cells (#CRL-1573, ATCC) were grown in Dulbecco’s Modified Eagle Medium (DMEM; #SH30022.01, Cytiva, Marlborough, MA) supplemented with 10% fetal bovine serum (FBS) and 100 units/mL penicillin-streptomycin (PS) in a humidified incubator with 5% CO_2_ at 37°C. FreeStyle 293-F cells (#R79007293F, ThermoFisher) were grown in FreeStyle 293 Expression Medium (#12338026, ThermoFisher). Cells were cultured in shaking flasks on an orbital shaker platform at 130 rpm in a humidified incubator with 8% CO_2_ at 37°C. The cells were maintained at a low density of 0.2-2 million/mL.

#### CuFi-8 cells

Human primary airway epithelial cells isolated from a cystic fibrosis patient, were immortalized by the expression of human telomerase reverse transcriptase (hTERT) and human papillomavirus (HPV) E6/E7 oncogenes (49). These cells were cultured on collagen-coated 100-mm dishes or 6 well plates using PneumaCult-Ex Plus medium (#05040; StemCell Technologies, Vancouver, BC).

### Human airway epithelium cultured at an air-liquid interface (HAE-ALI)

Proliferating CuFi-8 cells dissociated from flasks and loaded onto Transwell permeable supports (#3470; Costar, Corning, NY) at a density of 1.5 × 10^5^ cells per insert with PneumaCult-Ex Plus medium in both the apical and basal chambers. 2 to 3 days after seeding, the media were replaced with PneumaCult-ALI medium (#05001; StemCell) in the basolateral chamber only. The cells were then differentiated/polarized in PneumaCult-ALI medium at an air-liquid interface (ALI) for 3–4 weeks (48). The maturation of the polarized HAE-ALI cultures derived from CuFi-8 cells was determined by measuring transepithelial electrical resistance (TEER) with a Millicell ERS-2 volt-ohm meter (MilliporeSigma, St Louis, MO). ALI cultures with a TEER value of >1,000 Ω·cm² were used for experiments.

### Neuraminidase (NA) treatment

NA (#11585886001) was purchased from MilliporeSigma (St. Louis, MO), and was reconstituted in double-distilled water to a final concentration of 5 U/mL, following the manufacturer’s instructions. For neuraminidase treatment, HEK293 cells or HAE-ALI cultures were washed 2-3 times with Dulbecco’s Phosphate Buffered Saline (DPBS; #SH30028.03, Cytiva), and then treated with NA at 50 mU/mL (100 mU/mL for HAE-ALI) in DMEM medium without FBS and PS at 37°C, 5% CO_2_, for 2 hours. After incubation, the cells were washed once with DPBS.

### Plasmids

#### rAAV production plasmids

Plasmids pAAVR2C5, pAAVRep2Cap2.5T (pR2C2.5T), and pAV2F5tg83luc-CMVmCherry (4.6-kb), and pHelper have been described previously (38).

#### Lentiviral vector

Guide(gRNA)-expressing lentiviral vectors for gene KO were constructed by inserting the targeting sequences of single guide (sg)RNAs into lentiCRISPRv2 (#52961, Addgene, Watertown, MA). The targeting sequences for *SLC35A1* and *TMED10* KO are 5’-ATA AAG TTA TTG CTA AGT GT-3’ and 5’-TCT AGG ATC ACG AGT TGG TC-3’, respectively. For *KIAA0319L* and *TM9SF2* KO, we used the sequences previously described (38).

#### SLC35A1 expression plasmids

SLC35A1 ORF and the mutants, T128A and ΔC Tail, were codon-optimized and synthesized at Twist Biosciences (South San Francisco, CA). They were cloned in pLenti CMV Blast empty (#17486, Addgene).

### Lentivirus production and transduction

#### Lentivirus production

Lentiviruses were produced by transfecting HEK293T cells with sgRNA-expressing plentiCRISPRv2 plasmids, along with two packaging plasmids, psPAX2 and pMD2.G, using PEI MAX as described previously (45). The lentiviruses were then concentrated through a 20% sucrose gradient by ultracentrifugation in a SureSpin 630 rotor (Thermo Scientific) at 19,400 rpm for 3 hours. The transduction units of the produced lentiviruses were titrated using qPCR Lentivirus Titer Kit (#LV900, ABM, Richmond, BC, Canada).

#### Lentivirus transduction

HEK293T cells were transduced at a multiplicity of infection (MOI) of 5 transduction units per cell. Two days post-transduction, the cells were treated with puromycin at a final concentration of 2 µg/mL to select for the pool of transduced cells or single-cell colony expansion.

#### rAAV production

rAAV5, AAV2, and AAV2.5T vectors were produced by triple transfection of HEK293 cells with plasmids encoding the rep and cap genes, the adenoviral helper functions, and the vector genome containing the transgene. Vectors were purified using CsCl gradient ultracentrifugation followed by dialysis against PBS (56,57). The purified vectors were quantified by quantitative (q)PCR using a mCherry-specific probe as previously described (57). rAAV1, 3, 4, 6-9, and 11-13 were purchased from AAVnerGene (Rockville, MD).

### gRNA library and genome-wide CRISPR/Cas9 screen

A genome wide CRISPR/Cas9 screen was conducted in 293-F cells. The Brunello CRISPR/Cas9 knockout library was utilized for the genome-wide screen. Cells were transduced with lentiviral vector lentiCas9-Blast (#52962-LV, Addgene) and selected with blasticidin. Cas9-expressing cells (blasticidin-resistant) were then transduced with the Brunello lentiCRISPR gRNA library (#73178-LV, Addgene) and selected with puromycin. The double-resistant cells were expanded for rAAV5 transduction, followed by flow cytometry (FACSArialll, BD Biosciences, San Jose, CA) to collect the mCherry-negative cells. After two rounds of selection, the genomic DNA (gDNA) from sorted cells was extracted for next-generation sequencing (NGS).

### gDNA extraction, NGS, and bioinformatics analysis

#### gDNA extraction

The cells of the unsorted control (gDNA^Sort0^), the first (gDNA^Sort1^) and the second (gDNA^Sort2^) sorted groups were subjected to extraction of gDNA using the Blood and Cell Culture DNA Midi Kit (#13343; QIAGEN, Germantown, MD).

#### NGS

The gDNA samples were subjected to PCR-based amplification of guide sequences and indexed according to the protocol from the Broad Institute of MIT and Harvard (58). The PCR amplicons were sequenced using the Illumina NextSeq 2000 platform.

#### Bioinformatics analysis

NGS data were analyzed using the MAGeCK software package for sgRNA recognition sequences (42). Significance values were determined after normalization to the control population, and the data were reported as -log_10_ (Enrichment score). The analyzed data were visualized using Prism 10 (GraphPad). Genes were categorized by gene ontology (GO) terms using PANTHER v19.0 (59,60). The hits, represented by the enrichment score, were plotted along the y-axis and arbitrarily scattered within their categories along the x-axis. The size of the dot was determined according to the fold changes between sorted and unsorted groups.

### CRISPR/Cas9-based gene KO

HEK293 cells and CuFi-8 cells were transduced with lentiviral vectors expressing target-specific gRNAs. Transduced cells were selected with puromycin, and the KO efficiency was confirmed by Western blotting. The gene KO CuFi-8 cells were then differentiated at an air-liquid interface for polarized airway epithelial cultures (HAE-ALI).

### rAAV Transduction

For monolayer cultured cells, the cells were seeded overnight in 48-well plates. rAAV was added to each well at a multiplicity of infection (MOI) of 20,000 DNase digestion-resistant particles (DRP)/cell. The transduction efficiency was analyzed using a firefly luciferase assay 3 days post-transduction.

For the transduction of HAE-ALI, 100 µL of DPBS-diluted rAAV2.5T was added to the apical chamber of the transwell at an MOI of 20,000 DRP/cell. Subsequently, 0.5 mL of culture media was added to the basolateral chamber. After 16 hours, all liquid in the apical and basolateral chambers was removed and the chambers were washed three times with DPBS, pH 7.4 (Corning). Fresh culture media were then added to the basolateral chamber.

### Firefly luciferase assay

Gene knockout cells were transduced with rAAV vectors carrying a firefly luciferase reporter gene. Transduction efficiency was measured by quantifying luciferase activity using the Luciferase Assay System (Promega, Madison, WI) according to the manufacturer’s instructions. Luminescence was measured with Synergy LX Reader (BioTek, Santa Clara, CA).

### Vector binding and entry Assay

For binding assays, cells were incubated with rAAV vectors at 4°C for 2 hours. Unbound virions were removed by washing with PBS, and cells were lysed for qPCR quantification of bound vector genomes. For entry assays, cells were incubated with rAAV at 37°C for 2 hours, washed, and treated with trypsin to remove surface-bound virions. Cells were then lysed, and internalized vector genomes were quantified by qPCR.

### Cell fractionation

Cell fractionation was performed using the Subcellular Protein Fractionation Kit (#78840, ThermoFisher) (38). Cytoplasmic and nuclear fractions were isolated from transduced cells, and vector genomes in each fraction were quantified by qPCR.

### Vector genome quantification

Total DNA was extracted from cells or the cell-subfractions using the DNeasy Blood & Tissue Kit (Qiagen, Hilden, Germany). Vector genomes (DRP) were quantified by qPCR using primers specific for the rAAV genome (transgene: *mCherry*) (57). Standard curves were generated using known amounts of vector DNA to calculate genome copy numbers.

### Immunofluorescence assay and confocal microscopy

Cells were fixed with 4% paraformaldehyde, permeabilized with 0.1% Triton X-100, and blocked with 5% BSA in PBS. Cells were then incubated with primary antibodies followed by fluorescently conjugated secondary antibodies. Nuclei were stained with DAPI. Images were captured using a confocal microscope (CSU-W1 SoRa, Nikon, Melville, NY).

### Lectin and staining

Biotinylated *Sambucus nigra* lectin (SNA)(#B-1305), biotinylated *Maackia amurensis* lectin II (MAL II) (#B-1265), and fluorescein-conjugated *Erythrina cristagalli* lectin (ECL) (#FL-1141) were purchased from Vector Laboratories (Newark, CA).

Cells were fixed with 4% paraformaldehyde and blocked with carbo-free blocking solution (#SP-5040-125, Vector Laboratories). Cells were then incubated with biotinylated lectins followed by DyLight 649-conjugated streptavidin (#SA-5649-1, Vector Laboratories). Then the cells were permeabilized with 0.1% Triton X-100, and the nuclei were stained with DAPI. Images were captured using a confocal microscope Leica SP8 STED (Leica Microsystems, Deerfield, IL) or CSU-W1 SoRa (Nikon, Melville, NY).

### Flow cytometry

Flow cytometry was carried out as previously described (61). Briefly, the treated cells were washed twice with DPBS, dissociated using Accutase, and blocked with carbo-free blocking solution (#SP-5040-125, Vector Laboratories). The cells were then incubated with a biotinylated lectin (at 1:500 dilution in DPBS) for 30 min on ice, followed by staining with fluorescein isothiocyanate (FITC) conjugated streptavidin (#SA-5001-1, Vector Laboratories) for 15 min on ice. The cells were analyzed on a 5-laser spectral flow (Aurora; Cytek Biosciences, Seattle, WA), and data were analyzed using FlowJo v10 software (FlowJo, LLC, Ashland, OR).

### Sodium dodecyl-sulfate polyacrylamide gel electrophoresis (SDS-PAGE) and Western blotting

Cells were collected and lysed as previously described (62,63). The lysates were separated by SDS-PAGE and transferred to PVDF membranes. Membranes were blocked with 5% non-fat milk and incubated with primary antibodies followed by infrared dye-conjugated IgG (H+L) secondary antibody. Finally, the membrane was imaged on a LI-COR Odyssey imager (LI-COR Biosciences, Lincoln, NB).

### Antibodies used in this study

#### Primary antibodies

Rabbit anti-SLC35A1 (#A10658), rabbit anti-TMED10(#A18090), and mouse anti-β actin (#AC004) were purchased from ABclonal (Woburn, MA). Rabbit anti-KIAA0319L (#21016-1-AP) was obtained from Proteintech (Rosemont, IL). Sheep Anti-TGN46 (#GTX74290) was purchased from GeneTex (Irvine, CA), rabbit anti-TM9SF2 (#95189) was sourced from NOVUS (Centennial, CO). Anti-AAV5/2.5T intact particles (#03-651148) was purchased from ARP American Research Products (Waltham, MA).

#### Secondary antibodies

Alexa Fluor 488-conjugated donkey anti-sheep IgG (H+L) cross-adsorbed secondary antibody (# A-11015), Alexa Fluor 647-conjugated goat anti-mouse IgG (H+L) cross-adsorbed secondary antibody (# A-21235), and Alexa Fluor 594-conjugated goat anti-rabbit IgG (H+L) cross-adsorbed secondary antibody were purchased from ThermoFisher Scientific. DyLight 800-conjugated anti-rabbit IgG (#5151S) and DyLight 800-conjugated anti-mouse IgG (#5257S) were purchased from Cell Signaling (Danvers, MA).

### Statistical analysis

All data are presented as mean ± standard deviation (SD) obtained from at least three independent experiments by using GraphPad Prism 10. Statistical significance (P value) was determined by using an unpaired Student’s t-test for two groups or one-way ANOVA with post-hoc Bonferroni test for the comparison among more than two groups. ****P < 0.0001, ***P < 0.001, **P < 0.01, and *P < 0.05 were considered statistically significant, and NS represents statistically no significance.

## ACKNOWLEDGMENTS

The study was supported by NIH grants AI150877, AI156448, AI166293, AI180416, and HL174593, and Cystic Fibrosis Foundation grant YAN23G0. We are indebted to Dr. Richard Hastings at the Flow Cytometry Core Laboratory of the University of Kansas Medical Center, which is sponsored, in part, by the NIH/NIGMS COBRE grant P30 GM103326 and the NIH/NCI Cancer Center grant P30 CA168524. The super resolution confocal microscope Nikon CSU-W1 SoRa and Leica SP8 STED were supported by NIH S10 OD 032207 and NIH S10 OD 023625, respectively. The funder had no role in study design, data collection and interpretation, or the decision to submit the work for publication.

## DATA AVAILABILITY

All data used to evaluate the conclusions in this study are presented in the paper and/or the supplemental material. NGS data are available from the National Center for Biotechnology Information Sequencing Read Archive (SRA) under accession numbers SRR30593935 (Sort 0), SRR30594236 (Sort 2), SRR30594237 (Sort 1), and BioProject under accession number PRJNA1158467.

---

**Figure S1.**
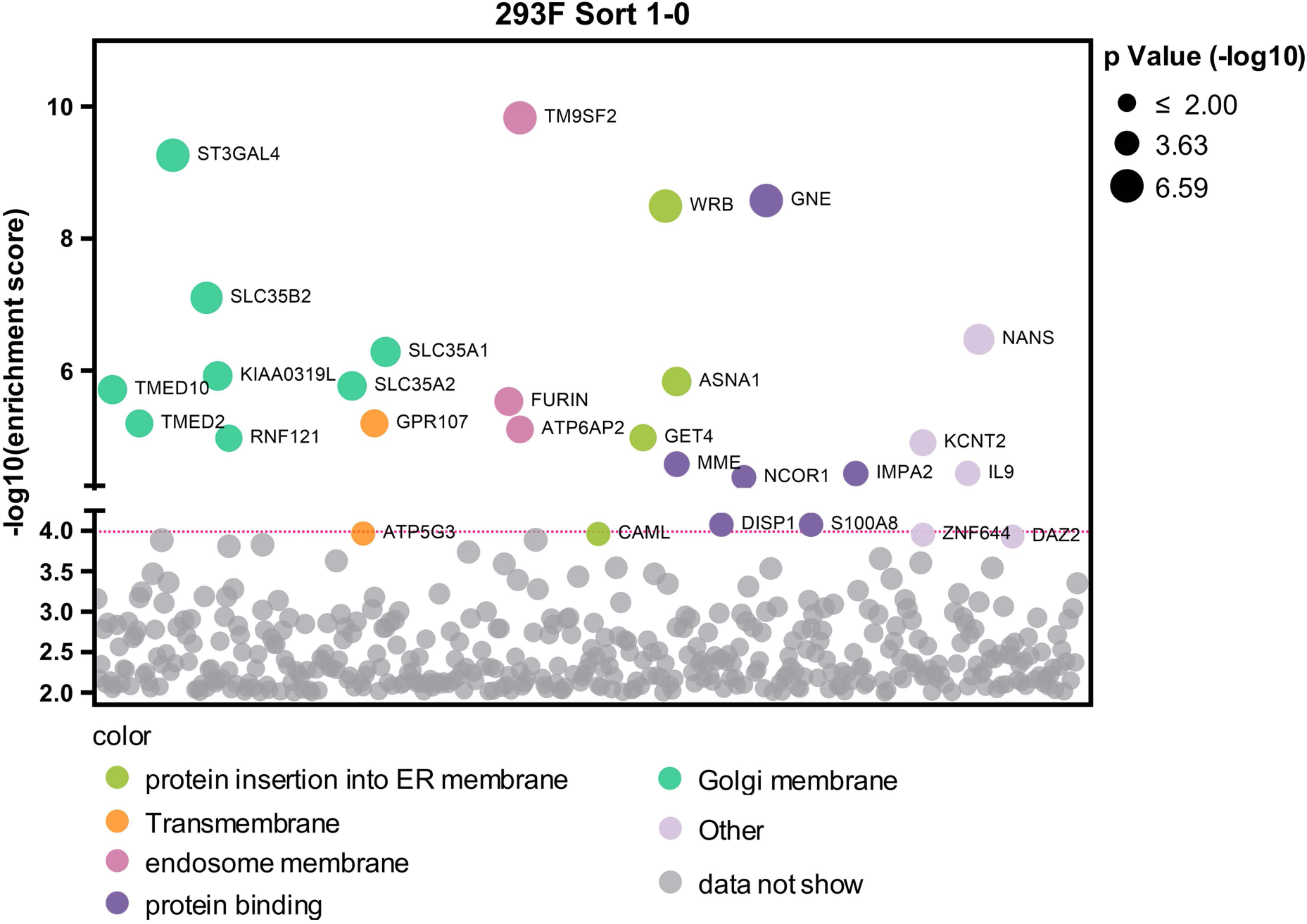
Genes enriched in the first-round screen of mCherry-negative cells. The x-axis represents genes targeted by the Brunello library, grouped by GO analysis. The y-axis shows the enrichment score [-log_10_] of each gene based on MAGeCK analysis of the sgRNA reads in gDNA^Sort1^ vs gDNA^sort0^ (**Table S2**). Each circle represents a gene, with its size indicating the statistical significance [-log_10_] of enrichment when comparing gDNA^Sort1^ to gDNA^Sort0^. The color of each circle represents the function of the genes. Only genes with an enrichment score greater than 10^4^ are shown.

**Figure S2.**
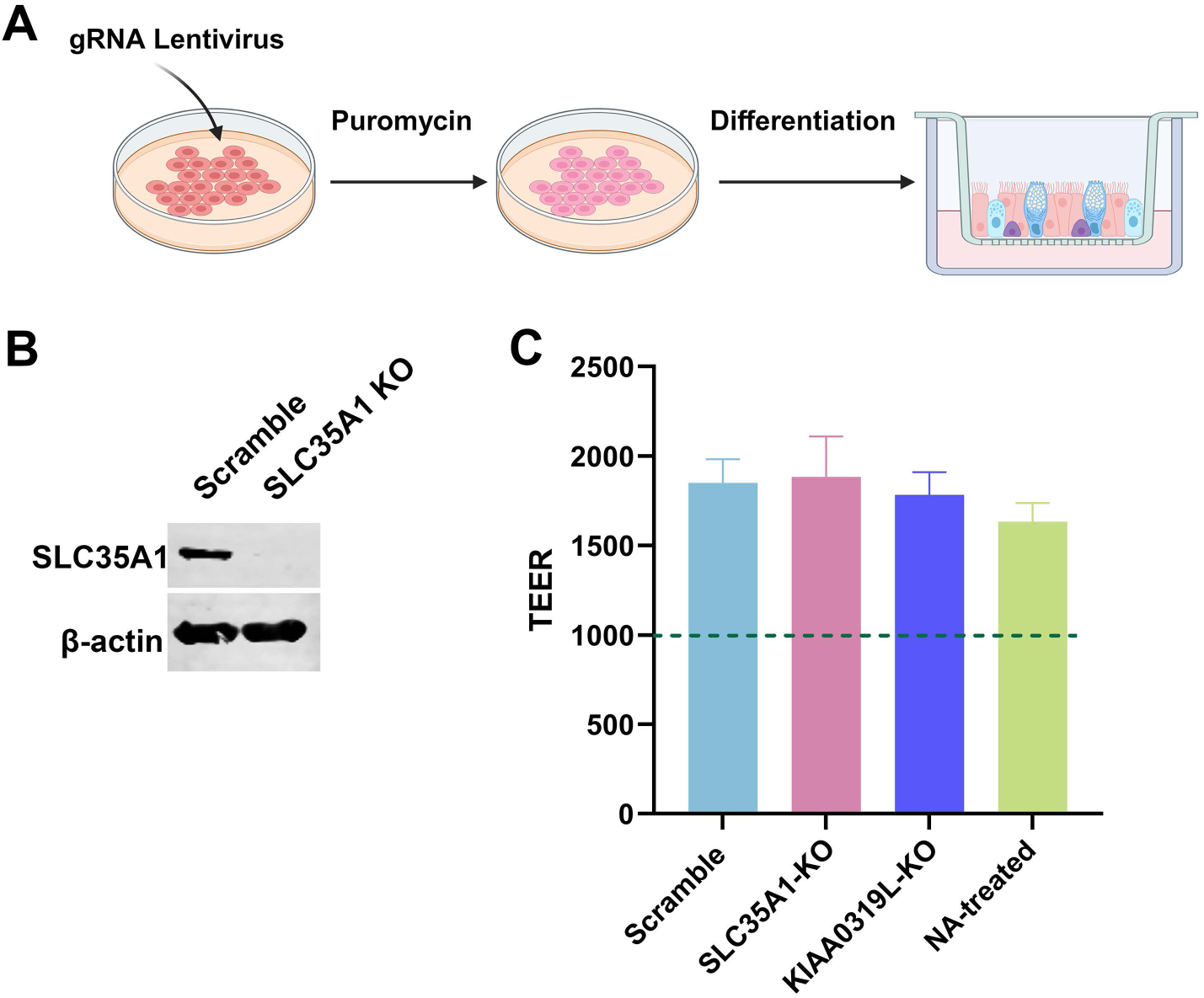
SLC35A1 KO in HAE-ALI culture. **(A) Generation of HAE-ALI^SLC35A1-KO^ cultures.** Human airway epithelial cell line CuFi-8 cells were transduced with a gRNA/Cas9 lentivirus. The puromycin resistant cells were seeded onto Transwell inserts and differentiated at an ALI for 4 weeks. **(B) Validation of SLC35A1 expression in HAE-ALI cultures.** Western blotting detected SLC35A1 expression in cultures derived from the scramble control but none in the cultures from *SLC35A1* KO cells. β-actin was detected as a loading control. **(C) Transepithelial electrical resistance (TEER) measurement.** HAE-ALI cultures, Scramble control, SLC35A1-KO, KIAA0319L-KO, and the Scramble control treated with NA were detected for TEER values.

**Figure S3.**
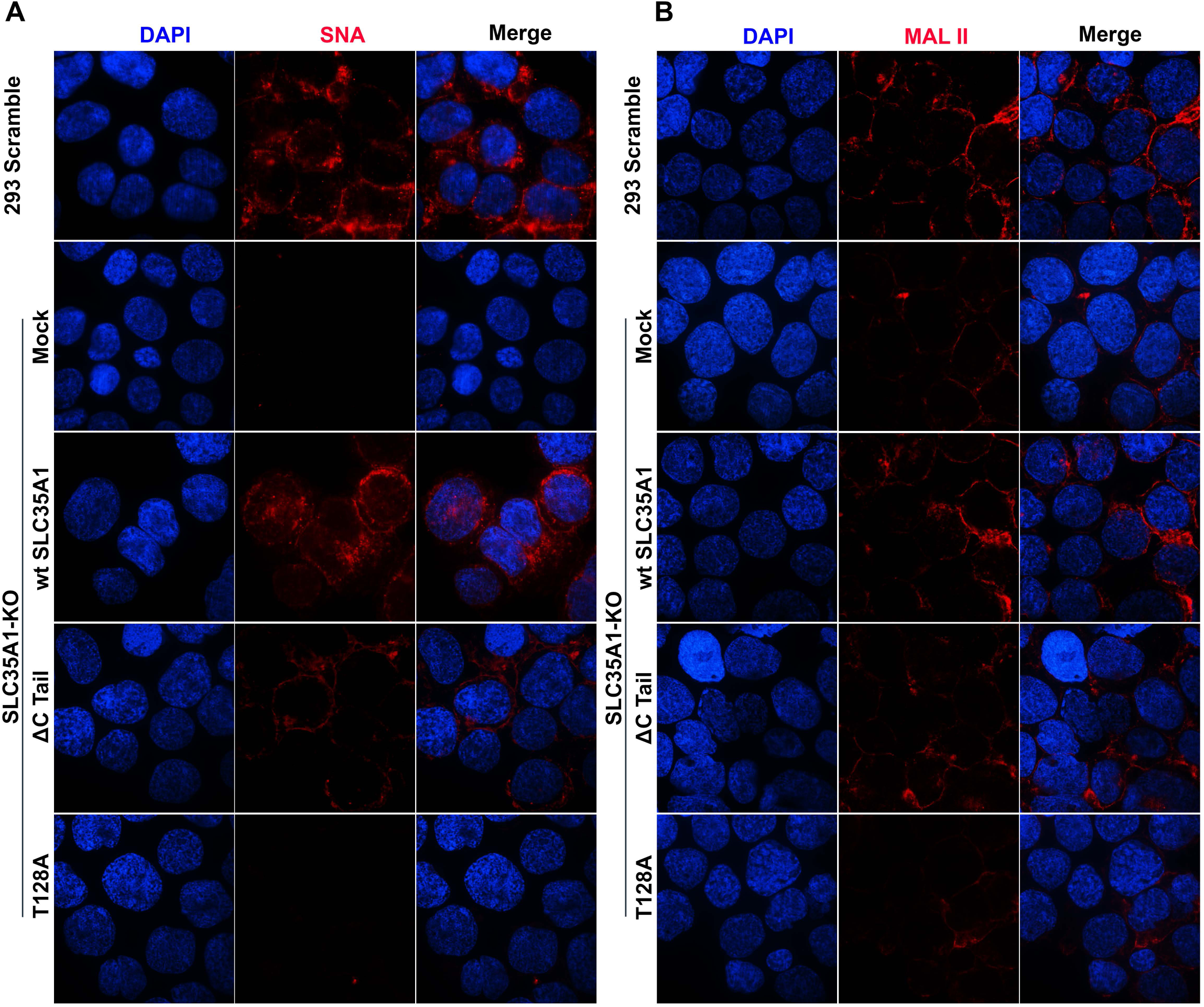
SIA expression in wide-type or mutant *SLC35A1* expression HEK293^SLC35A1-KO^ cells. HEK293^SLC35A1-KO^ cells were mock-treated or transduced with lentiviral vectors expressing SLC35A1 WT, T128A and ΔC Tail mutants, as indicated, followed by selection of blasticidin (at 10 µg/ml) for 2 weeks. The cells were fixed with 4% PFA and then permeabilized with 0.1% Trixon X-100 for intracellular staining. Biotinylated SNA and MAL II lectins were used to stain glycan expression in HEK293^SLC35A1-KO^ cells. SNA (**A**) and MAL II (**B**) stained cells were incubated with DyLight 649-conjugated streptavidin for visualization under a confocal microscope (CSU-W1 SoRa, Nikon) at 60×. HEK293 Scramble cells were used as a control.

**Table S1. A list of genes enriched in the second round (FD400210-FD400208) of the sorted mCherry-negative cells and ranked by the -log_10_ enrichment score.**

**Table S2. A list of genes enriched in the first round (FD400209-FD400208) of the sorted mCherry-negative cells and ranked by the -log_10_ enrichment score.**

